# Flow Physics Explains Morphological Diversity of Ciliated Organs

**DOI:** 10.1101/2023.02.12.528181

**Authors:** Feng Ling, Tara Essock-Burns, Margaret McFall-Ngai, Kakani Katija, Janna C. Nawroth, Eva Kanso

## Abstract

Organs that pump fluids by the coordinated beat of motile cilia through the lumen are integral to animal physiology. Such organs include the human airways, brain ventricles, and reproductive tracts. Although cilia organization and duct morphology vary drastically in the animal kingdom, ducts are typically classified as either carpet or flame designs. The reason behind this dichotomy and how duct design relates to fluid pumping remain unclear. Here, we demonstrate that two structural parameters – lumen diameter and cilia-to-lumen ratio – organize the observed duct diversity into a continuous spectrum that connects carpets to flames across all animal phyla. Using a unified fluid model, we show that carpet and flame designs maximize flow rate and pressure generation, respectively. We propose that convergence of ciliated organ designs follows functional constraints rather than phylogenetic distance, along with universal design rules for ciliary pumps.

## Main

To perform their physiological functions, many organs in animal biology rely on luminal flows driven by the coordinated beat of motile cilia. In humans, ciliated ducts pump fluids in the airways [1], brain ventricles and spinal canal [2], and reproductive system [3]. Their failure is directly linked to major pathologies, including bronchiectasis [4], hydrocephalus [5], and ectopic pregnancy [6]. Cilia in these ducts are usually short relative to the lumen diameter and are oriented perpendicular to the epithelial surface, in a ciliary carpet design [7]. Many animals also feature ducts with a strikingly different cilia arrangement, the ciliary flame design [8], where tightly packed, comparatively long cilia beat longitudinally along a narrow lumen. Ciliary flames that are thought to pump fluid for the purpose of excretion provide a model system for studying human kidney disease [9, 10].

Despite the fundamental importance of ciliated ducts in animal physiology, functional studies of intact ciliated ducts are rare [11, 12], and the relationship between ciliated duct morphology and their ability to pump fluid remains largely unexplored. Ciliary flames and carpets have not been compared functionally, nor have other ciliated duct morphologies been characterized in relation to these two fundamental designs.

Existing studies have focused on exposed ciliated surfaces and externally ciliated organisms because it is difficult to measure ciliary beat and fluid flow in intact internal ducts. Cilia oscillations, metachronal coordination, and microscale flows have been observed in microdissected *ex-vivo* epithelia [13], engineered *in-vitro* tissues [14, 15], and, more recently, in organ-on-chip models [16, 17]. Additionally, leveraging the remarkable conservation of the ultrastructure of motile cilia among eukaryotes, diverse protist and animal model systems have emerged for probing the functional spectrum of cilia, from locomotion and feeding [18, 19] to symbiotic host-microbe partnerships [20, 21]. External cilia that beat longitudinally in solitary or pairwise configurations, such as in mammalian sperm cells or in the unicellular protozoa *Euglena* and algae *Chlamydomonas*, are often called eukaryotic flagella and drive motility and gait transition in these microorganisms [19, 22–24]. But when longitudinally-beating cilia occur internally in a bundle, such as in planarians, they are labeled as ciliary flames, and are thought to be associated with filtration [8–10].

Despite remarkable progress in understanding the functional diversity of cilia, including the physical mechanisms underlying these functions [25–29], to date, the architecture of intact ciliated ducts inside of organisms is yet to be related to their pumping performance. Establishing this connection would elucidate the apparent dichotomy between carpets and flames, inform our understanding of human ciliopathies [30], aid in comparing different internal fluid transport mechanisms in animals [31, 32], and help in assessing the hypothesized roles of ciliated ducts in excretion [9, 10] and host-microbe interactions [33].

In this study, we leverage two giant larvaceans (*Bathochordaeus stygius* and *Bathochordaeus mcnutti*) as powerful model systems to study intact ciliated ducts. Larvaceans belong to Tunicata (Urochordata), a sister clade of vertebrates, and are an emergent model system for the study of gene regulation, chordate evolution, and development [34]. Giant larvaceans feature a multitude of ciliated ducts with both carpet and flame designs with exceptional optical access [35, 36], providing uniquely ideal conditions for studying the fundamental properties of cilia-generated flows in intact ciliated ducts with direct relevance to other chordates including humans.

To derive universal structure-function relationships, we combined our functional studies in giant larvaceans with an exhaustive review of published ciliated ducts in animals, and with physics-based computational models. Specifically, we (i) studied the larvacean system to characterize the morphological features and fluid transport mechanisms in flames versus carpets. We then used these insights to (ii) conduct a morphometric analysis of ciliated ducts across all animal phyla and identify a two-dimensional (2D) morphospace that organizes the design of ciliated ducts by fluid transport function, and to (iii) explain this structure-function relationship through a unifying physics-based model that links duct morphology to fluid pumping in terms of flow rate and pressure generation. We arrive at a functional assessment of ciliated ducts based on morphology that could aid in studying their role in human disease as well as inspire engineering applications [37–39].

### Comparison of ciliary carpet and flame designs

Microscopic inspection revealed that most ciliated ducts in giant larvaceans, such as in the pharynx, esophagus and gut, exhibit the ciliary carpet design associated with mucus clearance and fluid circulation in humans [40] (Fig. 1A). The ciliary flame design, which is associated with filtration in planarians [9], was found in the giant larvaceans’ prominent ciliated funnel (Fig. 1A and Extended Data Fig. 1), an organ whose function remains obscure [41]. To compare and contrast carpet and flame designs, we measured their structural properties, ciliary activity, and fluid transport velocity in the intact animal. In the carpet-style esophagus, cilia are much shorter than the width of the lumen (ca. 6 *µ*m vs. 100 − 200 *µ*m, respectively) (Fig. 1B) and they beat at ciliary beat frequencies (CBF) of 20 Hz with a stroke cycle perpendicular to the duct walls (Fig. 1D). Neighboring cilia coordinate their beat longitudinally along the duct and propel luminal fluid and food particles at flow speeds of up to 50 *µ*m s^−1^ (Fig. 1C, Supplementary Video 1). The ciliary beat frequency and flow parameters are comparable to those of other ciliary carpets, such as the mammalian airway (Extended Data Fig. 2A) and ependymal epithelia [2, 42].

**Figure 1.**
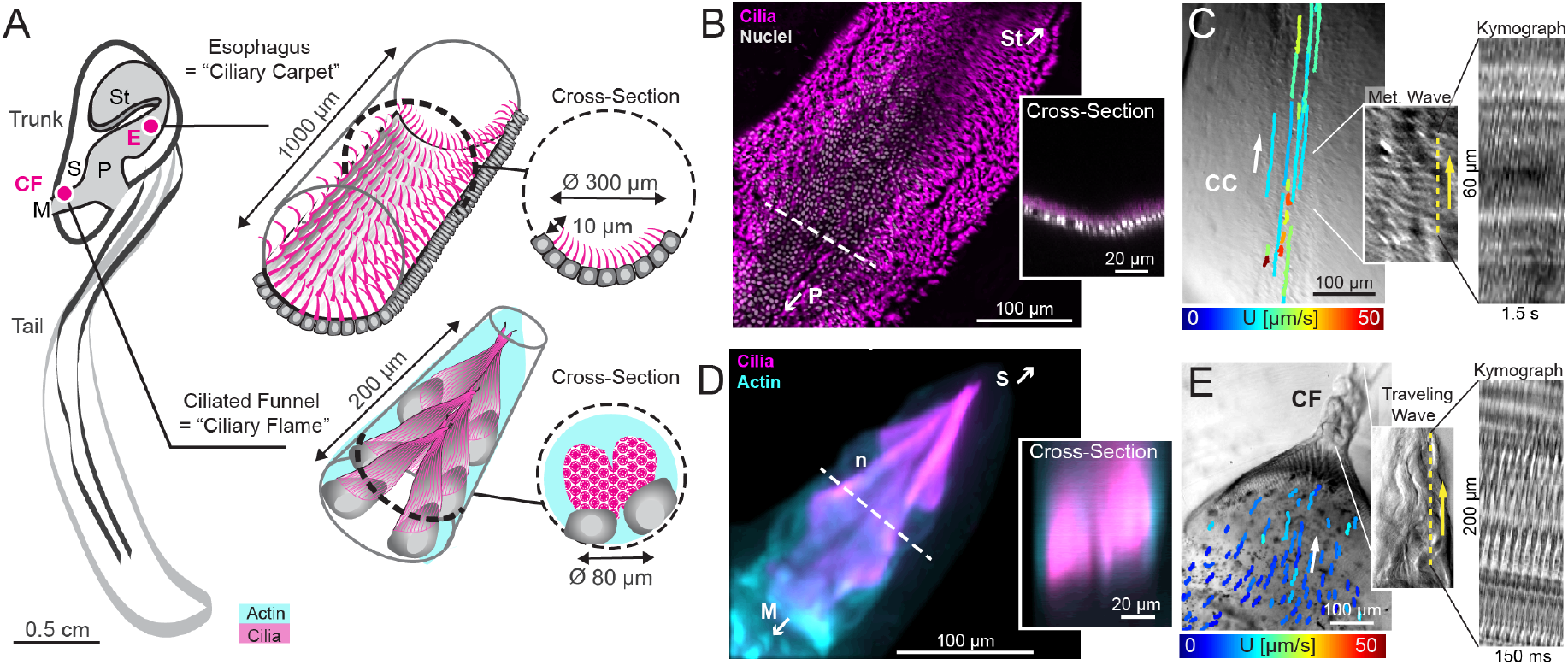
Comparison of ciliated ducts with ciliary flame and ciliary carpet designs in *B. mcnutti*. **A**. The stomach (St), pharynx (P), and esophagus (E) are lined with ciliary carpets (CC). A ciliated funnel (CF) containing ciliary flame cells connects the mouth (M) cavity to the internal blood sinus (S). Zoomed-in schematics show perspective and cross sectional view of ciliary carpets and funnel. **B**. Confocal image of the ciliary carpet of the esophagus (magenta: acetylated *α*-tubulin). Inset: Horizontal cross section at dotted line. The cilia only fill a small fraction of the lumen. **C**. Direction and magnitude of fluid flow driven by the esophageal ciliary carpet. Insets: Snapshot and kymograph of the esophageal carpet showing metachronal wave and beat frequency of ∼20 Hz. Yellow dash line shows the slice where kymograph is taken. Yellow arrow indicates direction of metachronal wave. **D**. Confocal image of the ciliary flame inside the ciliated funnel (magenta: acetylated *α*-tubulin); n, nucleus. Inset: Horizontal cross section at dotted line. **E**. Direction and speed of fluid flow converging into the ciliated funnel driven by the ciliary flame. Insets: Snapshot and kymograph of the ciliary flame showing traveling wave and beat frequency of ∼60 Hz. Yellow dash line shows the slice where kymograph is taken. Yellow arrow indicates direction of traveling wave. A-E: Similar results were confirmed in a total of three animals.

In contrast to the carpet-style esophagus, cilia in the larvacean funnel are longer than the width of the duct lumen (ca. 100 *µ*m versus 55 *µ*m, respectively) and align parallel to the lumen in a densely packed fashion (Fig. 1E and S.1B-D), which is typical of ciliary flames associated with filtration [8, 10]. While the beat kinematics of most ciliary flame systems are unknown, our data suggest that the coordinated motion of the many parallel cilia create periodic waves that travel along the entire length of the 100 *µ*m long larvacean flame (Supplementary Video 2). These cilia beat at a CBF of up to 60 Hz (Fig. 1g), and generate flow speeds of ca. 40 *µ*m s^−1^ towards the internal blood sinus (Fig. 1F, Supplementary Video 3). The kinematics of the larvacean ciliary flame are similar to those of the better studied, 20-times shorter ciliary flames of planarians (Extended Data Fig. 2B) [43], and appear to be a key feature by which ciliary flames transfer a directional force to the surrounding fluid regardless of flame size.

The stark structural differences between ciliary carpets and flames and the apparent lack of other duct designs in larvaceans underscore the questions of whether all ciliated ducts in nature fall into one of these two design categories and what the functional consequences are.

### Functional constraints of duct designs

To address these questions we performed an exhaustive survey on the presence, morphology, and reported fluid transport functions of ciliated ducts in all 34 animal phyla [44, 45]. Since kinematic data of ciliary beat was unavailable in most cases, we collected morphometric data accessible from still images of ciliated ducts, namely, cilia orientation relative to duct (perpendicular or longitudinal), duct length *L*, lumen diameter *H*, cilia length *h/*2, and cilia-to-lumen ratio *h/H*, which ranges from 0 to 1 and represent the relative fraction of the duct taken up by cilia, independent of cilia orientation (Fig. 2A and Extended Data Fig. 3). We collected these data from published work and our own imaging analysis of multiple species of Urochordates and Mollusks (Extended Data Fig. 4), covering a total of 61 duct systems representing 26 animal phyla (Supplementary Table 1). The remaining 8 phyla either lacked evidence of motile cilia or ciliated ducts or, in the case of sponges (phylum Porifera), were excluded because of their complex duct morphologies [46, 47].

**Figure 2.**
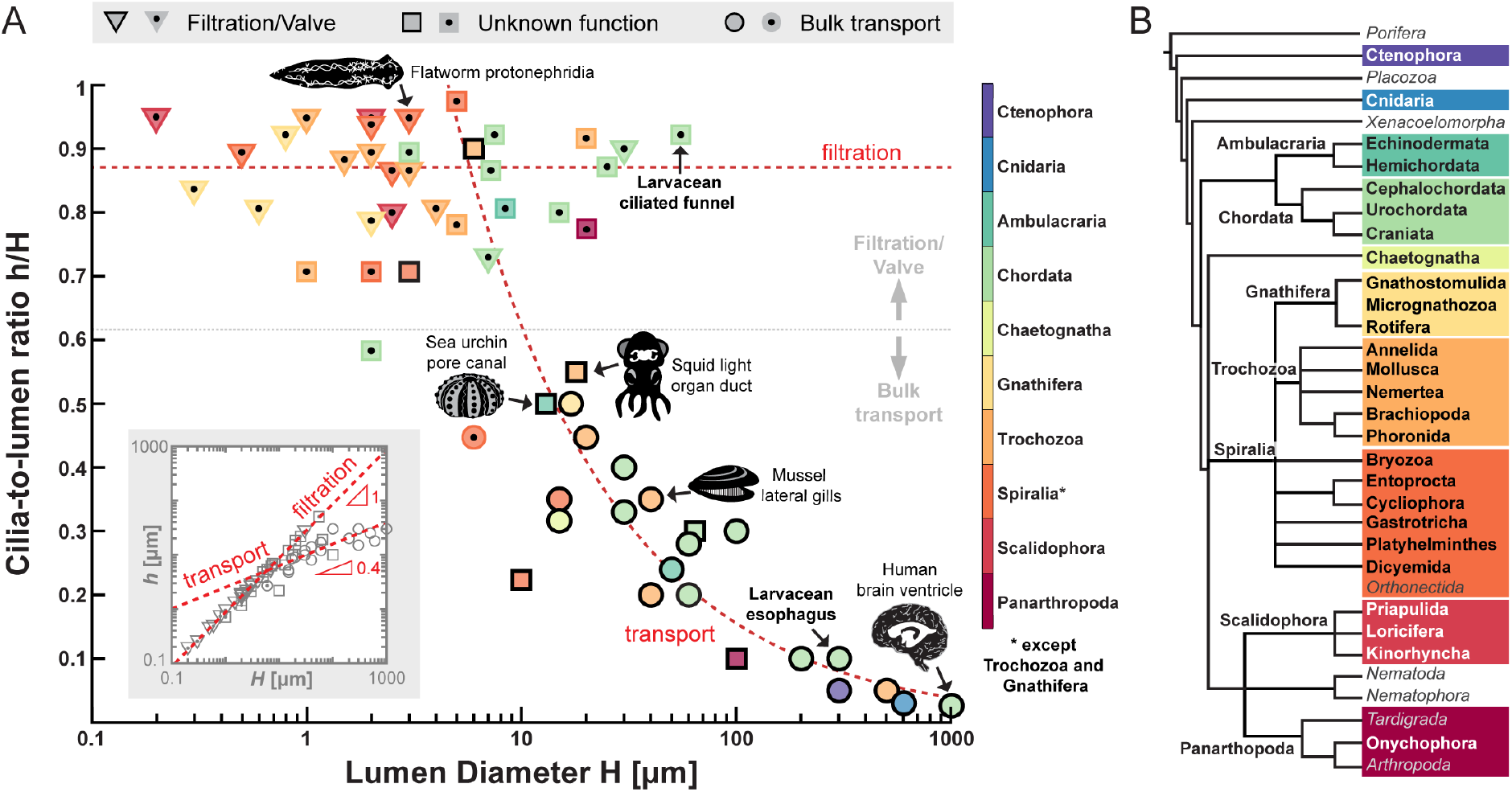
The morphospace of ciliated ducts in nature. **A**. Ciliated ducts plotted based on cilia-to-lumen ratio *h/H* and lumen diameter *H* and classified by their functions and phyla (see Extended Data Fig. 5). Specific duct examples are illustrated by schematics. Triangles ▽ represent systems whose primary functions are thought to be (ultra-) filtration, excretion, or pressure generation. Circles ∘ denote systems that transport or mix food particles, liquids, or cells. Squares □represent systems whose functions are currently unknown or disputed. Symbols with solid boundaries indicate perpendicular ciliation while symbols with a central dot have longitudinal ciliation. Symbol color indicates phylogenetic clades listed in the colorbar. Dotted grey line indicates the *h/H* value identified by machine learning to divide systems with filtration function from systems with bulk transport function. Power-law fits for systems with bulk transport and filtration functions are shown as red dashed lines, respectively, with their coefficients shown in the inset. **B**. Phylogenetic tree showing phyla known to feature ciliated ducts and included in the analysis (bold type) and the phyla where ciliated ducts are absent, not documented, or shaped in a way that cannot be captured by the morphospace in (A). The color code is the same as in (A). Tree design adapted from [44, 45]

Using a cross-validated machine learning approach, we probed the ability of the morphometric parameters to correctly predict whether a given ciliated duct was reported to perform bulk transport (clearance, circulation, or cargo transport) or serve as a filter/valve. We found that the cilia-to-lumen ratio *h/H* was a perfect predictor of known ciliated duct function and, when plotted against lumen diameter *H* to visualize the spatial scales involved, organized the 61 surveyed duct systems by function (Fig. 2B and Extended Data Fig. 5). A value of *h/H >*0.62 was associated with filtration or valve function, and a value of *h/H <*0.62 indicated bulk fluid transport Mapping cilia orientation onto the morphometric distribution further showed a clear association of filtration/valve function with flame designs (longitudinal cilia and high values of *h/H*) and of bulk transport with carpet designs (perpendicular cilia and low values of *h/H*). Cilia orientation by itself was not an optimal predictor of function due to marked exceptions, such as longitudinally aligned (but sparse) cilia in the collecting ducts of planarian excretory organs, which perform bulk transport of fluid [8]. Other morphometric parameters, e.g., duct length or diameter, were at best weakly predictive of ciliated duct function (Supplementary Fig. S1).

The *h/H* against *H* plot showed a continuous distribution of designs spanning carpets to flames, rather than a strict dichotomy. Ciliated duct designs can thus be described by their unique combination of cilia-to-lumen ratio and lumen diameter. In general, higher cilia-to-lumen ratios tended to associate with narrower ducts and flame designs, and lower cilia-to-lumen ratios with wider ducts and carpet designs, whereas systems with large *H* and large *h/H*, for example, were markedly absent. Importantly, the data associated with filtration and transport functions separated into two power laws: for transport (circles), *h* ∝ *H*^0.4^ while for filtration (triangles), *h* ∝ *H* (inset Fig. 2A). The cases with unknown function (squares) spanned both trends. Mapping the two power laws onto the morphospace (*h/H, H*) (dashed lines, Fig. 2A) emphasized the strong correlation between the fluid function – filtration or transport – and the ciliated duct morphology. Since the distribution of systems in the morphospace was not reflected by their phylogenetic distance but organized by their fluid transport activity (Fig. 2 and S.5), these data suggest a convergent evolution of ciliated duct designs based on function. Consistently, while the larvacean esophagus and ciliated funnel represent opposite ends of the morphospace (Fig. 2A), they belong to the same species.

Intuitively, the design of ciliated ducts for fluid pumping is constrained by the metabolic cost of building and actuating the cilia [48, 49], limiting the number and length of cilia on a given ciliated surface. A higher cilia-to-lumen ratio *h/H* also implies that less space in the duct lumen can be taken up by fluid. Minimizing the cost of cilia production along a cell surface and the cost of cilia activity while maximizing fluid volume seems to favor a low cilia-to-lumen ratio at large lumen diameter. However, the flame cells observed in purported filtration systems suggest that other functional considerations are at play. Specifically, structural data suggest that flame cells achieve filtration by pumping fluid through a flow-resisting sieve [8, 50, 51], indicating that higher cilia-to-lumen ratios may be specialized to pump fluid in the presence of high resistance to flow, or, equivalently, in the presence of *adverse pressure gradients* that cause counter-currents relative to the desired flow direction.

We hypothesized that lower cilia-to-lumen ratios which transport fluids at comparatively greater lumen diameters and flow rates, are able to pump flows at low or zero adverse pressures, whereas higher cilia-to-lumen ratios are limited to small diameters and low flow rates because of the high density of active ciliary material required to overcome adverse pressure gradients. These trade-offs would explain both the non-uniform distribution of ciliated duct designs and their functional classification.

### A unified model of fluid transport in ciliated duct designs

To rigorously probe this hypothesis, we used a Brinkman-Stokes model, where a cylindrical duct of diameter *H* is lined by a porous layer of overall thickness *h* representing the ciliary layer, enclosing an inner free lumen of diameter (*H* − *h*) (Fig. 3 and Supplementary Methods §.2). This lumped-layer model emulates ciliary carpets for *h* small relative to *H* and resembles ciliary flames as *h* approaches *H*. It thus bridges the two archetypes, carpets and flames, in a concise way that is unachievable using existing envelope [13, 52–56] and discrete cilia models [25, 57–63] and enables efficient exploration of the entire morphospace spanned by *h/H* and *H* independent of cilia orientation, including designs not (yet) known to biology.

**Figure 3.**
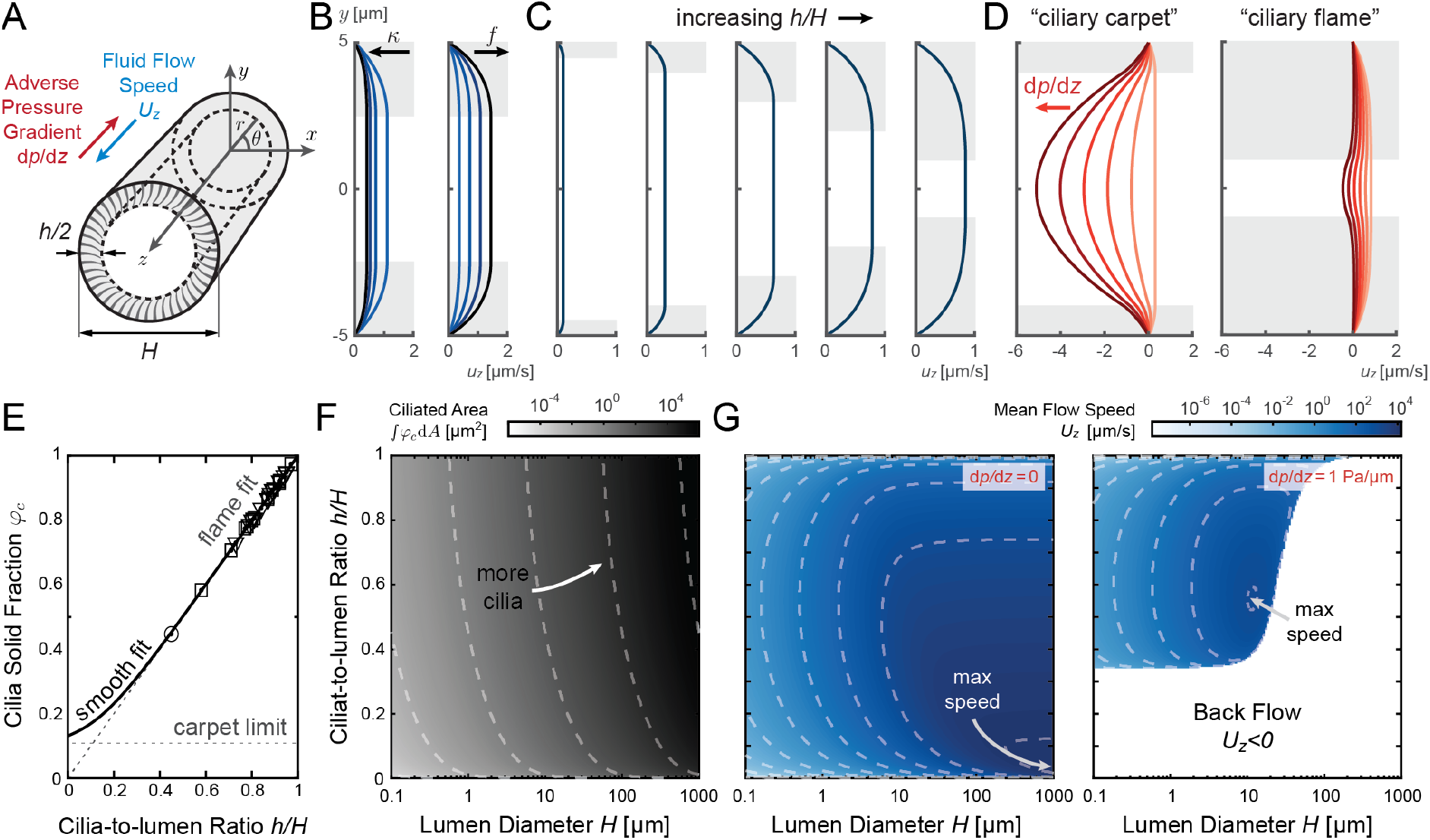
Brinkman-Stokes model maps ciliated duct morphology to fluid pumping. **A**. Schematic of axisymmetric two-layer model of ciliated duct: cilia are homogenized into an isotropic porous ring layer with height *h/*2, specific fluid fraction *φ* and cilia fraction *φ*_*c*_ = 1 − *φ*, Brinkman drag coefficient *K*_*c*_ = *κφ*_*c*_, and active force density *f*_*c*_ = *fφφ*_*c*_, all inside a duct with diameter *H*. **B**. The net flow *u*_*z*_ decreases with increasing resistance of the porous layer *κ* (here, *κ/µ* = 0.5, 1, 1.5, 2[*µ*m^−2^]), and increases with increasing the cilia activity *f* (shown for *f/µ* = 0.5, 1, 1.5, 2[(*µ*m *·* s)^−1^]). **C**. The net flow *u*_*z*_ approaches a bowing maximum as the cilia-to-lumen ratio *h/H* increases (shown for *h/H* = 0.1, 0.2, 0.4, 0.6, 0.8). **D**. Adverse pressure d*p/*d*z* drives a backward Poiseuille flow that scales quadratically with the size of the free lumen. (shown for d*p/*d*z* = 0, 0.2, 0.4, 0.6, 0.8, 1 *×* 10^−3^[*Pa/um*]). Systems with small *h/H* (ciliary carpets) are less effective in pumping against adverse pressure compared to those with large *h/H* (ciliary flames). **E**. Empirical soft-plus fit of cilia fraction *φ*_*c*_ as a function of the cilia-to-lumen ratio *h/H*: ciliary flame data (symbols follows definition in Fig. 2A, only ciliary flame data points are shown) suggest that *φ*_*c*_ = *h/H* for large *h/H*, while ciliary carpet data indicate that lim 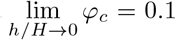. **F**. Total ciliated area over the entire morphospace (*H, h/H*): systems with higher *H* and *h/H* have more cilia per cross-section of the duct. **G**. Mean flow speed *U*_*z*_ at zero adverse pressure (left) and d*p/*d*z* = 1[*Pa/µm*] (right). In absence of adverse pressure, systems with small *h/H* and large *H* produces the fastest flow speed. Under adverse pressure, the maximum speed occurs at a larger *h/H* and smaller *H*. Dashed lines in F-G show the contour shape of the respective colormaps. Parameter values are set to: *H* = 10[*µm*] in panels B-D with light gray background indicating the ciliary layer. *φ*_*c*_ = 0.1 in B-D and, in F and G, it follows from the smooth fit of E. *κ/µ* = 1[*µ*m^−2^] in C-G, *f/µ* = 1[s *· µ*m^−1^] in C-D, *f* = 15[pN *· µ*m^−3^] in G, and *µ* = 10^−3^[Pa *·* s] throughout.

In the model, all characteristics of the ciliary layer were subsumed into three parameters: a cilia fraction *φ*_*c*_ and its complementary fluid fraction *φ* = 1 − *φ*_*c*_, a uniform force density *f*_*c*_ exerted by the cilia on the fluid (or the pressure gradient generated by ciliary force in a pipe [64]), and an effective Brinkman coefficient *K*_*c*_ encoding the drag resistance to fluid flows due to the presence of the cilia. This generic representation captures the two salient features of the ciliary layer – the cilia-generated force and drag-induced resistance to fluid flows – irrespective of the details of cilia distribution and beat kinematics.

In ciliary carpets, because cilia are mainly orthogonal to the duct surface, the cilia fraction *φ*_*c*_ can be estimated from longitudinal cross-sections of the duct, looking down at the density of cilia on the surface. Available images and our own data show little variations in *φ*_*c*_ across ciliary carpets for small *h/H*; we thus set 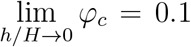 (Supplementary Methods §.3). In the flame systems, because cilia are mainly parallel to the duct wall, *φ*_*c*_ is determined by taking transverse cross-sections, and, by definition, is proportional to the cilia-to-lumen ratio *h/H* constructed from surveyed images. These observations inspire an empirical nonlinear fit of *φ*_*c*_ as a function of *h/H* that spans carpet and flame designs (Fig. 3E).

More cilia generate more force. By linearity of the dilute resistive-force theory in the Brinkman model, the ciliary force density *f*_*c*_ must scale linearly with the cilia fraction *φ*_*c*_. The fluid fraction *φ* is also important to transfer the ciliary force to the surrounding fluid; letting *f*_*c*_ be proportional to *φ* avoids the nonphysical situation of generating a ciliary force when *φ* = 0. We thus set *f*_*c*_ = *fφ*_*c*_*φ*, with a constant force density coefficient *f*. Meanwhile, more cilia induce higher resistance to fluid flows in the ciliary layer, and, by the same linearity argument, the effective Brinkman coefficient *K*_*c*_ = *κφ*_*c*_ must be proportional to *φ*_*c*_, where *κ* is a proportionality constant.

The downstream fluid velocities 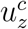 and 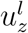 in the ciliary and free lumen layers are, by axisymmetry, function of the radial distance *r* only and are governed by the Brinkman and Stokes equations, respectively, with proper boundary conditions (BCs),

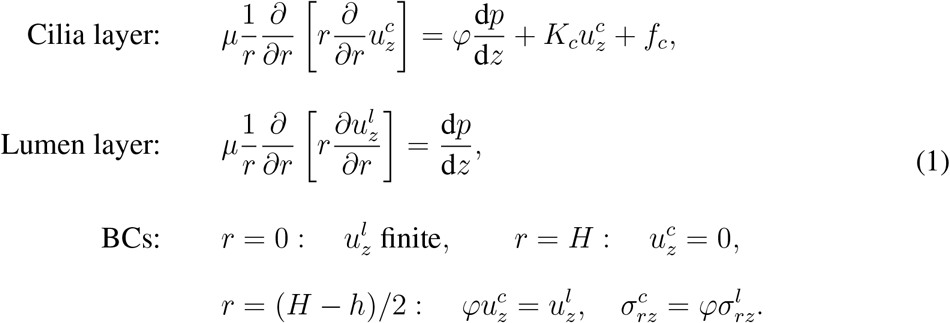

Here, *µ* denotes the fluid viscosity, d*p/*d*z* the adverse pressure gradient, and 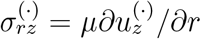 the fluid shear stress [65, 66]; see Supplementary Methods §.2for analytical solutions to (1).

For constant cilia fraction *φ*_*c*_, the profile of the fluid velocity *u*_*z*_(*r*), equal to *u*^*l*^ in the free lumen layer and *φu*^*c*^ in the ciliary layer, decreased with increasing Brinkman resistance *κ* and increased with increasing cilia forcing *f* and cilia-to-lumen ratio *h/H* (Fig. 3B-C). Importantly, in the presence of adverse pressure *dp/dz* ducts with low cilia-to-lumen ratio failed and allowed reversal of luminal flow, whereas positive luminal flow remained robust at high cilia-to-lumen ratio (Fig. 3D).

Considering the best fit of *φ*_*c*_ to experimental data (Fig. 3E), the corresponding ciliated area *A*_*c*_ =∫*φ*_*c*_d*A* varied over the morphospace (*H, h/H*) (Fig. 3F). We computed the associated net flows *U*_*z*_ = ∫ *u*_*z*_d*A* for all (*H, h/H*) at zero and non-zero adverse pressure (Fig. 3G). At *dP/dz* = 0, any ciliated duct design in the morphospace (*H, h/H*) can produce positive flow; however, speed decreased with increasing cilia-to-lumen ratio *h/H* for a given lumen diameter *H*.

This trade-off between maximizing net flows versus sustaining an adverse pressure is best appreciated in the pump function space (*Q*, d*p/*d*z*), where *Q* = ∫ *U*_*z*_d*A* is the net flow rate (Fig. 4A). Each ciliated duct operates along its own characteristic curve, connecting its maximal generated flow rate *Q* at d*p/*d*z* = 0 to the maximal adverse pressure d*p/*d*z* sustained without causing flow reversal *Q* = 0 as depicted in Fig. 4A, with solid pink lines corresponding to the (*H, h/H*) values of larvacean ducts in Fig. 1. Intuitively, a ciliated duct performs best when operating near the transition from maximal flow rate to maximal pressure.

**Figure 4.**
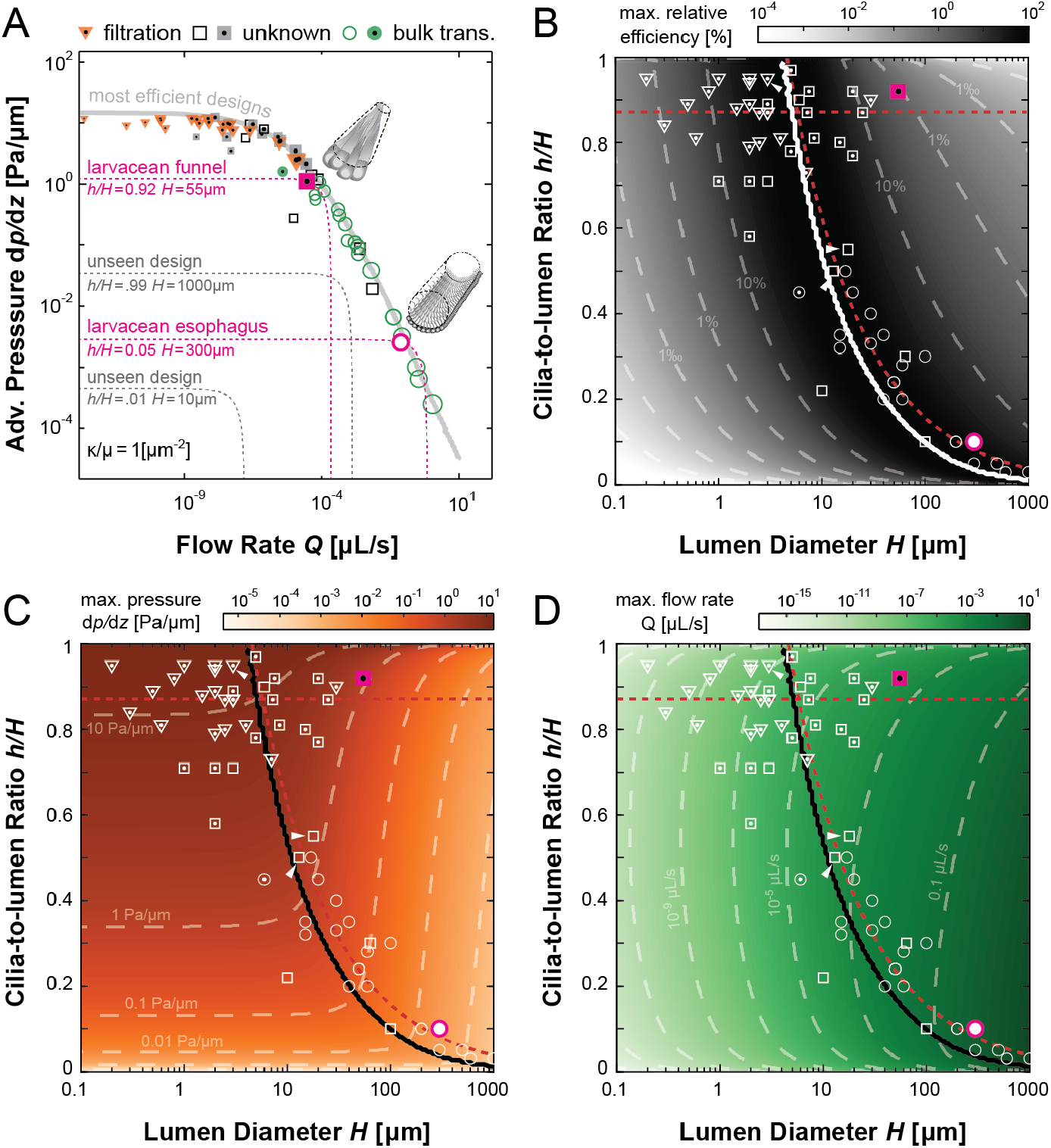
Optimal ciliated duct designs. **A**. Model prediction of pumping performance of optimal duct designs in the function space of flow rate vs. adverse pressure gradient (*Q*, d*p/*d*z*) are shown in the solid grey line. Non-optimal ducts (dashed grey lines) at small cilia-to-lumen ratio *h/H* and lumen diameter *H* generate lower pressure and flow rate, and non-optimal ducts at high *h/H* and *H* perform less than efficient while using more cilia. Here we fix the reference resistence constant *κ* to be equal to fluid viscosity *µ*. For results using other values of *κ* see Supplementary Fig. S2. Biological data from Fig. 2A are superimposed. Size of the markers indicates the relative amount of ciliated area per cross section (Fig. 3F). Operating curves (dashed pink lines) of the larvacean ciliated funnel and esophagus carpet are highlighted. Our model predicts that most biological ciliated ducts operate close to the optimal functional limits. **B**. Maximum relative efficiency (*E*_rel_ of Supplementary Alg. S1, solid white line) in the (*H, h/H*) morphospace, with data points of Fig. 2A (white symbols) superimposed. Most efficient designs continuously shift from carpet to flame designs. **C-D**. Maximum pressure (orange) and flow rate (green) generation in the (*H, h/H*) morphospace. Black solid lines are the morphologies with highest relative efficiency. Red dashed lines in B-D are the power-law fits shown in Fig. 2A, and gray dashed lines are the contour lines of the respective colormaps. *κ/µ* = 1[*µ*m^−2^], *f* = 15[pN*/µ*m^3^] with *µ* = 10^−3^[Pa *·* s] throughout.

### Optimal duct design depends on pressure requirements

To systematically probe optimal ciliated duct designs, we introduced a pumping efficiency *E* = *Q/A*_*c*_ defined as the ratio of the volumetric flow rate *Q* to total ciliary material *A*_*c*_ per cross sectional area. We sought optimal geometries (*H, h/H*) that maximize *E* under an imposed adverse pressure gradient d*p/*d*z*.

For a given d*p/*d*z*, we computed the flow speed *U*_*z*_ over the entire morphospace (*H, h/H*). At each (*H, h/H*), we calculated *Q* and *E* and identified the duct design (*H, h/H*) that maximized *E*. We assessed the performance of each (*H, h/H*) relative to the optimal design by computing its relative pumping efficiency *E*_rel_ scaled by the maximal pumping efficiency value.

We repeated this process for a range of d*p/*d*z* ∈ [10^−10^,10^2^] [Pa/*µ*m], and obtained the most efficient design values of (*H, h/H*) as a function of d*p/*d*z*; see also Supplementary Methods, Algorithm S.1.Optimization results are plotted on the function space (*Q*, d*p/*d*z*) (Fig. 4A, solid grey line). Results using the experimental values of (*H, h/H*) from Fig. 2A in our fluid model are superimposed (colored markers). The model predicts that the empirical ducts follow the optimal trend, with transport systems (circles) exhibiting larger *Q* values at lower d*p/*d*z* and filtrations systems (triangles) larger d*p/*d*z* at lower *Q*, and systems with unknown functions spanning both.

We plotted the optimal duct designs on the morphospace (*H, h/H*) (Fig. 4B, solid white line; for results using different values of *κ* see Supplementary Fig. S2) and depicted the highest relative efficiency *E*_rel_ as a colormap over the entire morphospace, which excludes designs both in the small *H*, low *h/H* and in the large *H*, high *h/H* corners of the morphospace. Designs that have relatively small lumen diameter *H* and low cilia-to-lumen ratio *h/H* are sub-optimal because for the same d*p/*d*z*, ducts with larger *H* are always more efficient in transporting flows (Fig. 4A, dotted grey lines). Designs that have relatively large *H* and high *h/H* are also sub-optimal because increasing the lumen diameter *H* limits the value of d*p/*d*z* that can be sustained at high *h/H*. Additionally, when increasing *H* at high *h/H*, the required ciliary material also grows, likely beyond biological limits.

Superimposing the biological data onto Fig. 4B, all surveyed designs aligned with the line that maximizes pumping efficiency (the median relative efficiency achieved is 57%; see Supplementary Fig. S3A). This line is also resembles the power-law fit for ducts with known bulk transport function (red dash line), with a slope of more than 0.3 in log-log scale (Supplementary Fig. S3B). Importantly, systems with presumed filtration functions are typically ciliary flames and occupy the region of the morphospace with smaller lumen diameter *H* and higher cilia-to-lumen ratio *h/H*. This zone is characterized by its ability to sustain higher maximum adverse pressure gradient just before flow reverses (*U*_*z*_ = *Q* = 0); see Fig. 4C. Conversely, systems whose primary function is to transport luminal fluid in bulk are typically ciliary carpets and thus reside in the region of the morphospace with larger lumen diameter *H* and lower cilia-to-lumen ratio *h/H*, a zone characterized by its ability to produce higher maximum flow rates (Fig.4D).

### Predicting pumping performance from structural features

We established a quantitative link between ciliated organ morphology and fluid pumping function. Our analyses suggest that the distribution of morphological features of ciliated ducts is not random; each biological system optimizes pump performance for specific pressure and flow rates. Systems with low cilia-to-lumen ratios maximize flow rate, while those with high ratios optimize pressure generation. Many systems exhibit intermediate ratios, likely enabling fluid transport against resistance. Our study also explains the absence of certain duct designs in our analysis due to inefficiency or biological limitations. Structural features of ciliated ducts can predict their ability to transport fluids or perform filtration. For example, our findings support the hypothesis that the larvacean’s ciliated funnel aids in fluid volume maintenance [67], rather than being limited to sensory and endocrine functions [41, 68, 69], and suggest that the Hawaiian bobtail squid’s ciliated conduit likely transports fluids in some areas and filters or pressurizes in others [33]. Dynamic changes in ciliated pore canals support a switch between fluid pump and valve functions in sea urchins [70].

Our findings shed light on waste excretion strategies in animals and may help explain why smaller animals rely on ciliary flame-based excretory organs while larger animals use muscle-powered hearts [71]. Assuming cilia have finite length and surface density, increasing lumen diameter eventually lowers the ratio of cilia to fluid volume, thus limiting the pressure and flow rates of individual flame pumps. Scaling up through the parallel operation of many small flame units would require a much larger organ volume than directly leveraging fluid pressures achieved by the heart (Supplementary Methods §.4). Even if cilia length was unlimited, advantageous scaling of energy efficiency in muscle versus ciliated cells could bias evolution towards muscle-powered filtration in larger animals.

Our analysis may inform disease phenotypes [72] and drug development targeting excretory functions of parasites [43]. Additionally, it offers insights for designing tissue-engineered and synthetic ciliary pumps and devices beyond carpet-like configurations [37–39, 73].

Limitations of our study include potential oversight of alternative duct designs in less studied animals and omission of not fitting our 2D morphospace, such as ciliated chambers in sponges [46, 47, 74] and ctenophores [75–78]. We did not differentiate between fluids or consider mixing flows, focusing on fundamental measures of pumping performance. Despite these limitations, our model provides a universal tool for assessing the physiological role of ciliated organs.

## Acknowledgements

This work was funded by NSF Inspire grant MCB1608744 (E.K. and M.M.-N.), NIH R01 HL153622 grant (E.K. and J.N.), ERC-STG grant 950219 (J.N.), NIH R37 AI50661, COBRE P20 GM125508, OD11024 and GM135254 grants (M.M.-N.), and David & Lucile Packard Foundation (K.K.). Acquisition of the Leica TCS SP8 X confocal microscope was supported by NSF DBI 1828262 (M.M.-N.).

## Authors contributions statement

Feng Ling: Partial Conceptualization, Investigation, Methodology, Formal Analysis, Software, Data curation, Validation, Visualization, Writing-Original draft preparation, Reviewing and Editing. Tara Essock-Burns: Investigation, Visualization, Writing - Reviewing and Editing. Margaret McFall-Ngai: Writing - Reviewing and Editing, Funding acquisition. Kakani Katija: Methodology, Investigation, Resources, Writing - Reviewing and Editing. Janna Nawroth: Conceptualization, Investigation, Methodology, Formal Analysis, Software, Data curation, Writing-Original draft preparation, Reviewing and Editing. Eva Kanso: Conceptualization, Investigation, Methodology, Formal Analysis, Writing-Original draft preparation, Reviewing and Editing, Supervision, Funding acquisition.

## Methods

### Ethical regulations

All applicable international, national, and/or institutional guidelines for the care and use of animals were followed. Animal collections were made on board RVs Western Flyer and Rachel Carson in the Monterey Bay National Marine Sanctuary (MBNMS). Activities were conducted under the MBARI institutional permit with MBNMS, specimen collections were performed in accordance with the guidelines of the California Department of Fish and Wildlife and collecting permits SC-200900003, SC-13337, and SC-190810004.

### Experimental methods

We collected four different larvacean species (*Bathochordaeus mcnutti, Bathochordaeus stygius, Fritillaria sp*., *Mesochordaeus erythrocephala*) and conducted in situ high-speed video microscopy of intact internal ciliated ducts to capture ciliary beat kinematics and fluid transport [20]. We also reanalyzed ciliary beat videos we previously recorded in human airway epithelial cultures [0]. To measure ciliated duct morphology, we employed immunofluorescence (IF) imaging of fixed samples of larvaceans and the Hawaiian bobtail squid (*Euprymna scolopes*). Additionally, we collected and analyzed duct morphology from micrographs in literature representing all animal phyla, exclusions applied. The morphometric data sets were analyzed using machine learning to identify the structural parameters most predictive of fluid transport functions (bulk transport or filtration/valving) reported in literature. Complete methods for data acquisition and analysis are provided in Supplementary Materials § S.1.

## Data availability

All source data is available in the manuscript or the supplementary materials.

## Code availability

All source code used to generate simulated data and figures is available in the manuscript or the supplementary materials.

**Extended Data Figure 1:**
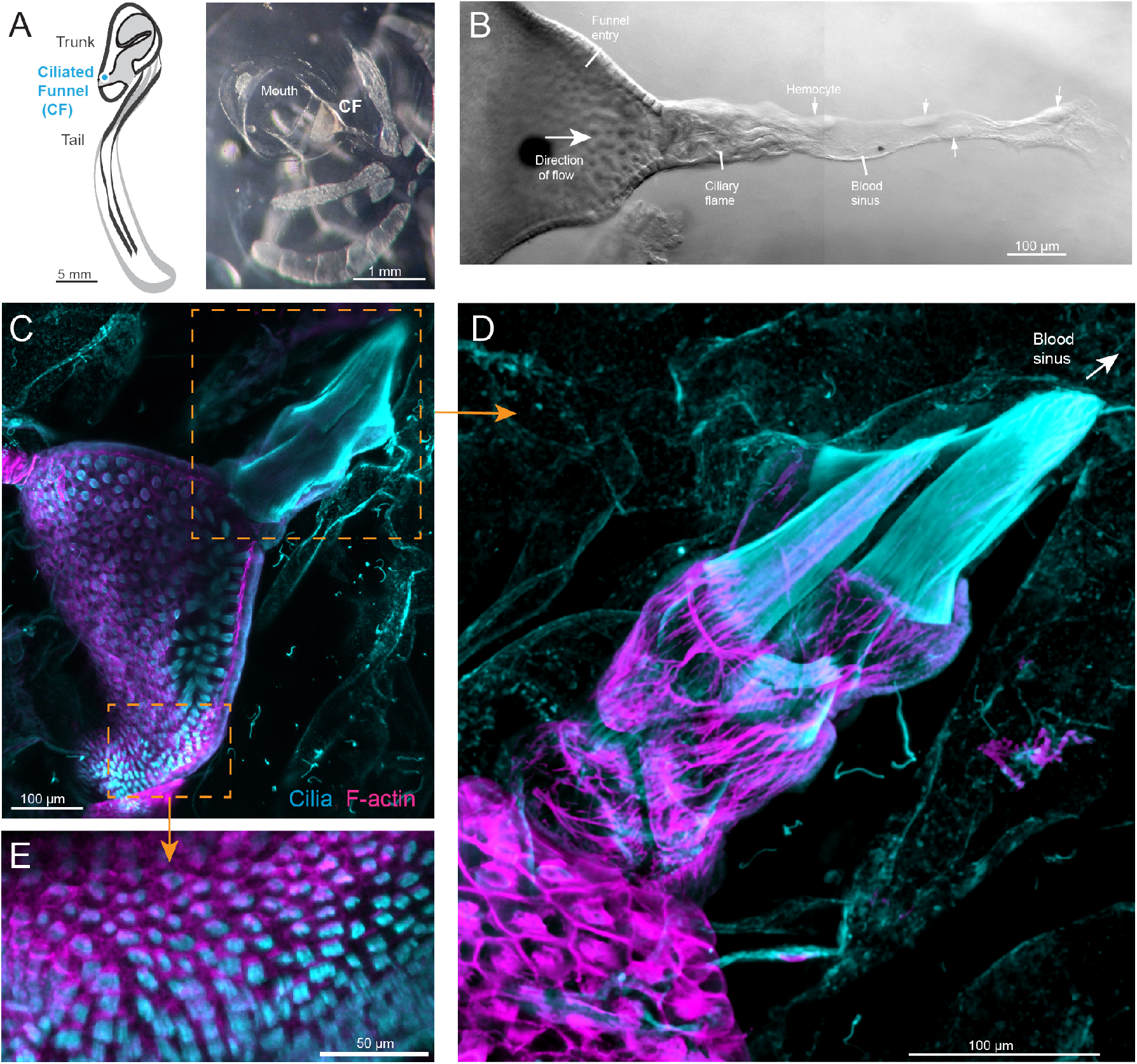
The flow generated by the ciliary flame of the giant larvacean appears to draw ambient seawater through a lattice of non-motile cilia, through the ciliary flame, and into the blood sinus. **A**, The ciliated funnel (CF) in the giant larvacean *B*.*stygius* branches off the mouth cavity (M). **B**, Phase-contrast cross-section of entire ciliated funnel shows protective lattice of non-motile cilia in the funnel entry (opening to mouth cavity), ciliary flame, and connection to blood sinus with putative hemocytes (small arrows). The direction of the cilia-driven flow is inwards (large arrow), consistent with a multi-stage filtration system [67]. **C**, Cross-sectional confocal image of the ciliated funnel in *B*.*stygius* including the funnel entry and the ciliary flame. **D**, Close-up on the ciliary flame, showing an actin mesh (magenta) encasing the large ciliary flame (cyan) which is composed of multiple, tightly packed ciliary flame cells and connects to the blood sinus. **E**, Close-up on the lattice of non-motile cilia that project into the funnel entry.The morphology shown in A-E was confirmed in a minimum of 3 animals.

**Extended Data Figure 2:**
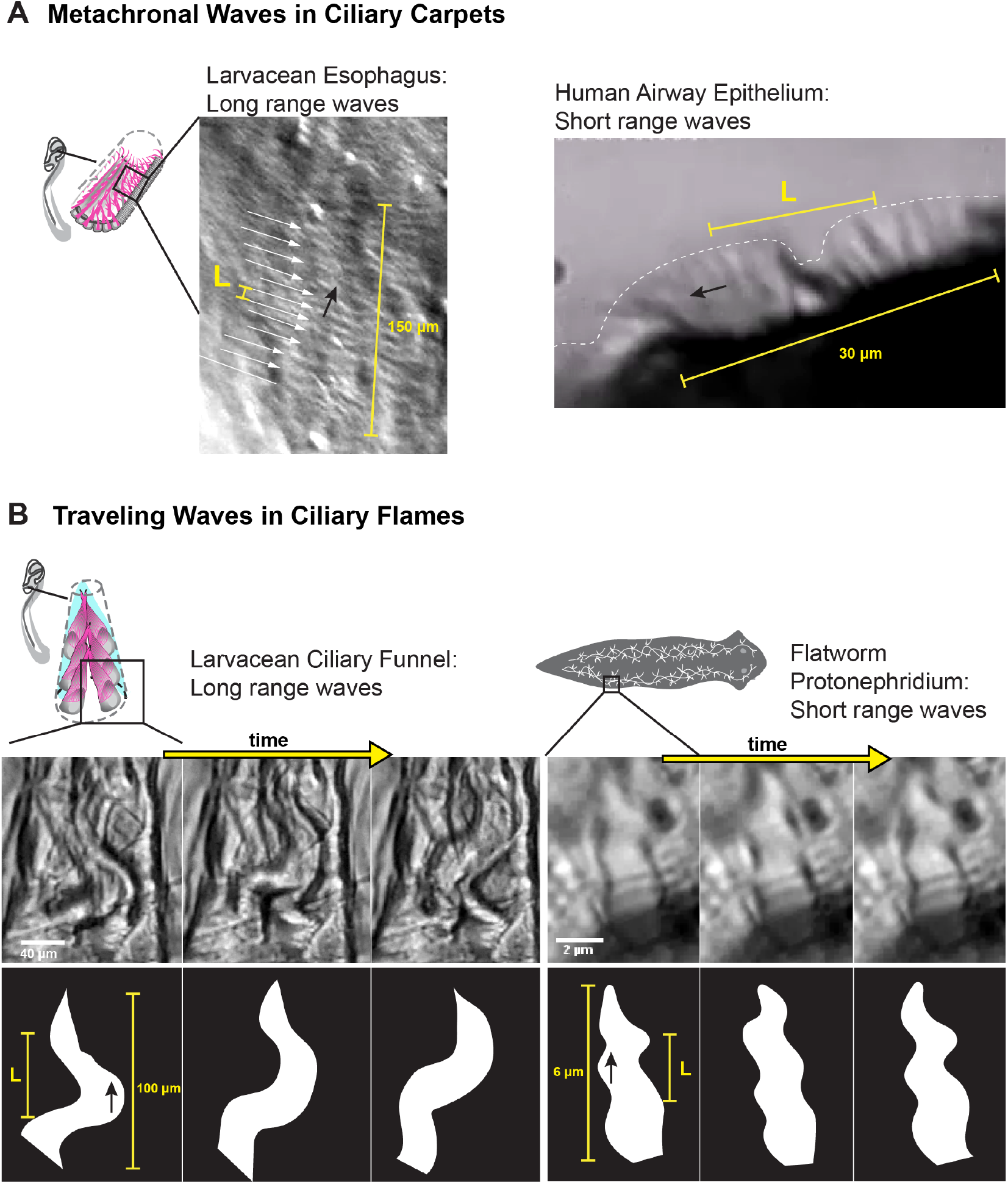
Ciliary beat coordination in metachronal and traveling waves. **A**, Ciliary carpets generate long-range or short-range metachronal waves of ciliary beat. Left, larvacean esophagus, arrows indicate crests of metachronal wave; right, engineered human airway epithelium. L, metachronal wave length. **B**, Ciliary flames generate long-range or short-range metachronal waves of ciliary beat. Left, time lapse of wave traveling along flame of larvacean ciliary funnel; right, time lapse of wave traveling along flame of planarian (flatworm) protonephridium. L, traveling wave length. The data shown in A-B was collected from one sample per species each.

**Extended Data Figure 3:**
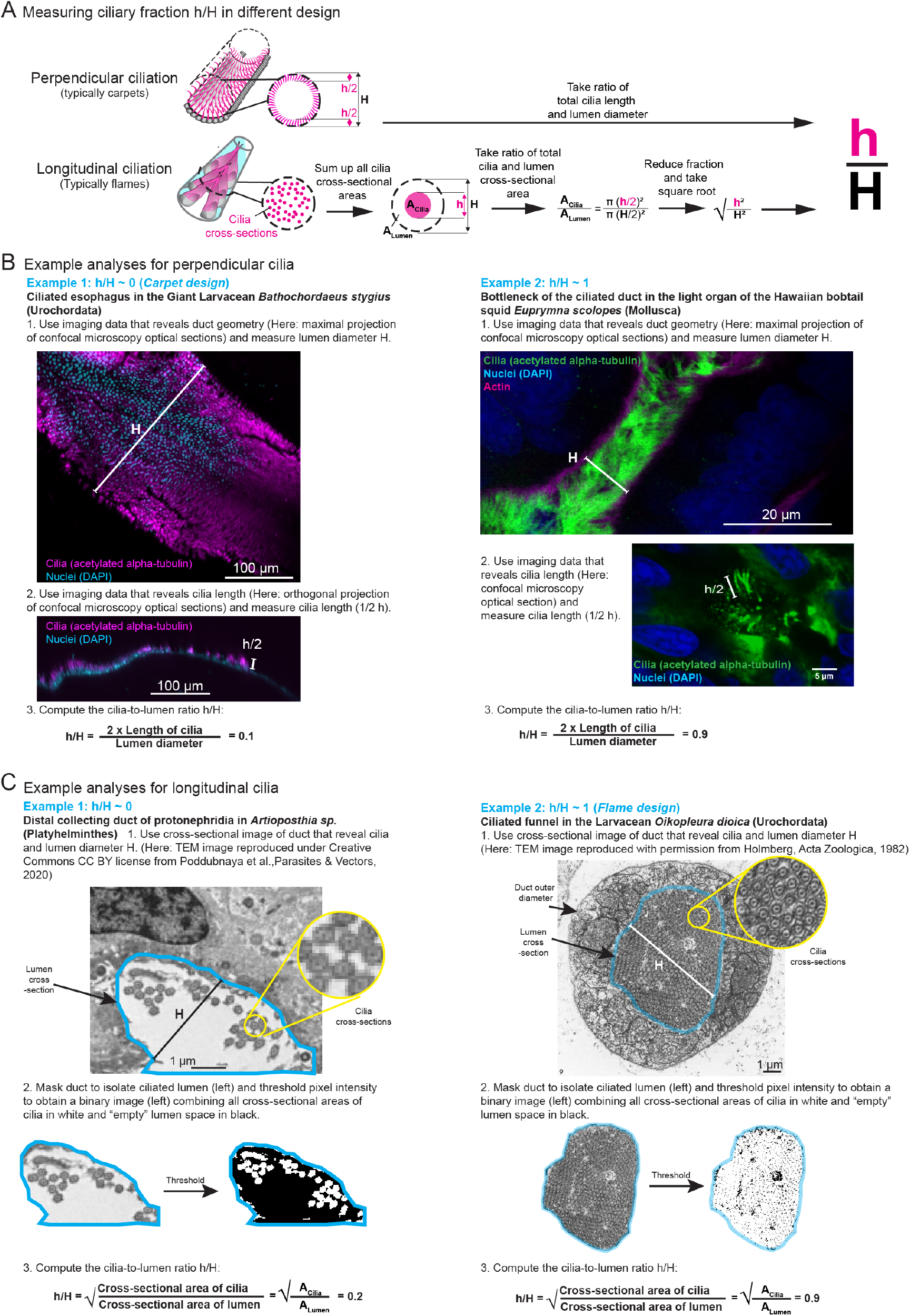
Methods to measure duct lumen (DL) diameter and cilia-to-lumen ratio. **A**,In carpet-style ciliated ducts *h/H* was determined as the ratio of the ciliary layer height h and the duct lumen diameter H. Since ciliary carpets are assumed to line both “floor”and “ceiling” of the ciliated duct, ciliary length corresponds to 1/2 h. In flame-style ciliated ducts the cilia are aligned longitudinally to the duct and hence cilia density rather than length determines the cilia-to-lumen ratio. h/H was therefore determined as the square root of the ratio of the cross-sectional area of the cilia to the cross-sectional area of the duct lumen. **B**, Example analyses of carpet designs with low *h/H* and high *h/H* values (own data from one sample). **C**, Example analyses of flame designs with low *h/H* and high h/H values. The left TEM image was adapted from [79] to highlight areas with sparse ciliation, under Creative Commons CC BY license. The right TEM image was adapted from [41], with permission from John Wiley and Sons.

**Extended Data Figure 4:**
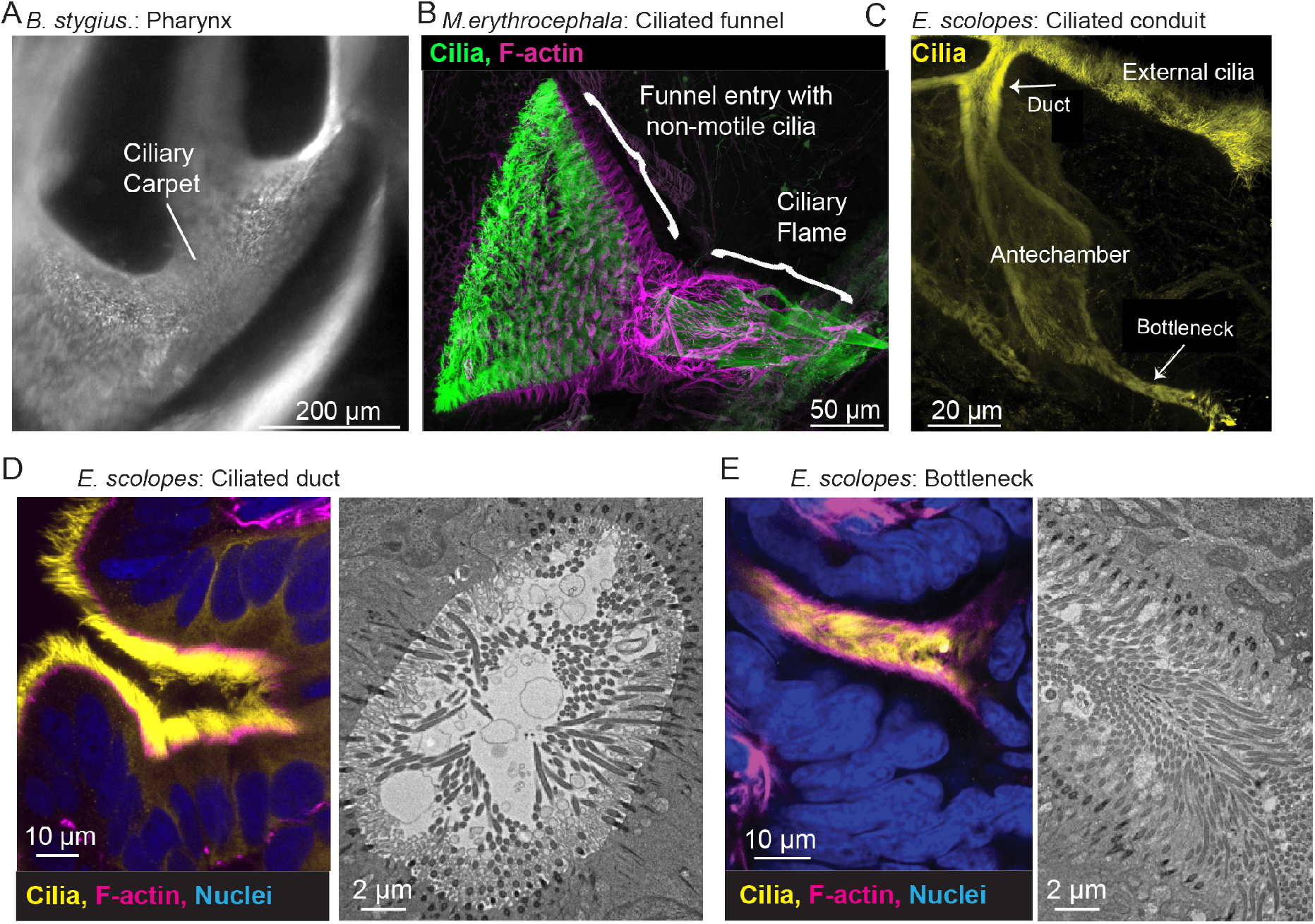
Carpet and flame-type ciliated ducts in Urochordates and Mollusks analysed in this study. **A**, The ciliated pharnyx in larvaceans, here *B. stygius* (Urochordata), is characterized by a ciliary carpet and a low h/H ratio. **B**, The ciliated funnel in larvaceans, here *Mesochordaeus erythrocephala* (Urochordata), is a flame design. **C**, The ciliated conduit of the light organ in the Hawaian Bobtail Squid *Euprymna scolopes* (Mollusca) features a carpet design in the duct and antechamber regions and a flame-like design in the bottleneck region, as seen in the close-up immunofluorescent and transmission electrode images of **D**, the ciliated duct and **E**, the bottleneck region. The data shown in A-B are taken in one animal each; data shown in C-E were validated in at least 3 animals as part of a published study [33]

**Extended Data Figure 5:**
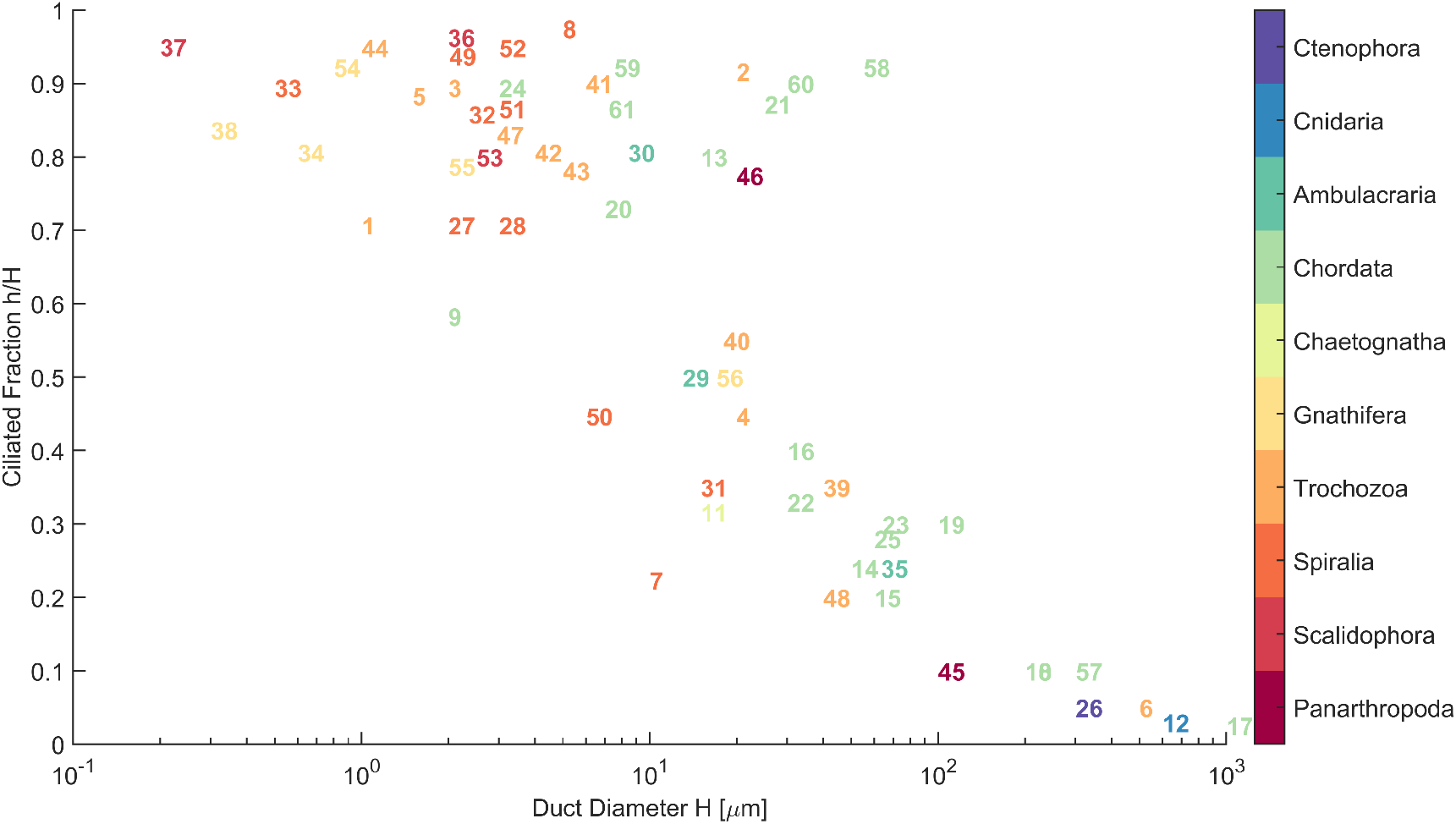
Indexed plot of cilia-to-lumen ratio as a function of lumen diameter. The plotted numbers correspond to the index numbers in Supplementary Table 1 and hence indicate the animal species corresponding to each data point, as well as the associated source of the data.

