## Supplementary Document for "Flow Physics Explains Morphological Diversity of Ciliated Organs"

### Supplementary materials

**Supplementary Video 1** *In vivo* beat kinematics and flow generation by the ciliated carpet of the esophagus in the giant larvacean. Video is slowed down 2x from original speed.

**Supplementary Video 2** *In vivo* beat kinematics of ciliary flame in the ciliated funnel of the giant larvacean. Note that video is slowed down 16x from original speed such that the flow moves very slowly.

**Supplementary Video 3** *In vivo* beat kinematics and flow generation of the ciliary flame in the giant larvacean. Video is slowed down 8x from original speed.

**Supplementary Table 1** Survey of structural and functional parameters of ciliated ducts in the animal kingdom (plus reference list of source publications).

**Supplementary Code** MATLAB code used to generate all simulated data in this manuscript.

#### S.1 Experimental Methods

##### S.1.1 Tissue Imaging

**Live imaging of larvaceans.** Four different larvacean species (*Bathochordaeus mcnutti, Bathochordaeus stygius, Fritillaria sp., Mesochordaeus erythrocephala*) were collected at water depths ranging from 55 m to 330 m in the Monterey Bay National Marine Sanctuary using acrylic detritus and suction samplers with remotely operated vehicles (ROVs) Ventana and Doc Ricketts as part of RVs Rachel Carson and Western Flyer cruises in May and October 2016, June 2018, and July 2021. Collected larvaceans were kept in seawater at 4°C for a maximum of 48 h before imaging analysis was conducted. For imaging, animals were placed in 10 cm petri dishes filled with seawater at 4°C that was regularly refreshed. The internal ciliated ducts were observed using phase contrast microscopy with 20x and 40x objectives. High-speed video microscopy was performed using either a Sony 4K Handycam FDR-AX700 or an SC1 camera (Edgertronic, CA, USA) mounted each with a custom optical adapter to the C-port or the eyepiece holder of the microscope. Ciliary beat frequency was measured from the videos using the kymograph plugin in ImageJ [1] which visualizes the number of beat cycles per second. Metachronal wave length and direction of propagation was derived from subsequent frames in high-speed video recordings as well as spacing and slope of the stripe pattern in kymographs. Flow was visualized by adding 1 to 10 $\mu$m sized glass microspheres microspheres (Dantec Dynamics, NY, USA) and 1 to 2 $\mu$m polymer microspheres (Cospheric, CA, USA) as tracer particles to the seawater. Fluid flow velocity magnitudes were measured from particle trajectories using the Trackmate plugin in ImageJ [2]. As the high density of cilia in the larvacean ciliated duct obscured the direct imaging of tracer particles, or prevented their passage, the fluid velocity magnitude within the duct was estimated from the flux in the funnel just outside the duct.

**Immunofluorescence (IF) imaging of larvaceans.** Animals were fixed in 4 percent paraformaldehyde in seawater for 24 h at 4°C, then washed 3x for 10-30 min in phosphate-buffered saline (PBS) and stored in PBS at 4°C until IF staining. For IF staining, the samples were incubated with primary monoclonal antibodies against $\alpha$-acetylated tubulin (to stain cilia; T6793, Sigma-Aldrich, MI, USA) in PBS for 24 h at room temperature, followed by 24 h of incubation in anti-mouse secondary antibody (Invitrogen, CA, USA), phalloidin (to stain F-actin), and DAPI (to stain nuclei) in PBS at room temperature. For IF imaging, tissues were mounted in a custom-made glass-bottom petridish and imaged with a Zeiss LSM 710 laser scanning confocal microscope using a 40x or 63x objective.

**IF imaging of *Euprymna scolopes*** Animals were cultured and samples were prepared and imaged using IF staining, laser scanning confocal microscopy, and transmission electron microscopy (TEM) as described previously [3].

#### S.1.2 Image Analysis

**Measurement of duct lumen diameter and cilia-to-lumen ratio.** Duct lumen (DL) diameter $H$ and cilia-to-lumen ratio $h/H$ were determined from own imaging data (Fig. 1, Extended Data Fig. 1 and 4) and micrographs found in literature (Supplementary Table 1). Two different methods were used for quantitative image analysis, depending on cilia orientation (Extended Data Fig. 3A). In perpendicularly ciliated ducts, which are typically carpets (Extended Data Fig. 3B, left) and only rarely present as highly occluded ducts (Extended Data Fig. 3B, right), $h/H$ was determined as the ciliary layer height $h$ divided by DL diameter $H$. To quantify $H$, we identified images that showed the duct in its full width or cross-section and directly measured the diameter. To quantify $h$, we identified images that showed close-ups of the ciliated carpet in cross-sectional or 3D perspective and we measured cilia length. Given $H$ and $h$, the cilia-to-lumen ratio $h/H$ is determined straightforwardly. Since perpendicularly ciliated ducts are assumed to line both "floor" and "ceiling" of the ciliated lumen, $h$ corresponds to twice the cilia length.

In longitudinally ciliated ducts, which are typically flames (Extended Data Fig. 3C, right) and only rarely feature sparsely ciliated designs (Extended Data Fig. 3C, left), the cilia are aligned longitudinally to the DL, and hence cilia density across the channel determines the cilia-to-lumen ratio, rather than cilia length. Thus, $h/H$ was determined from the summed cross-sectional area of all cilia divided by the total cross-sectional area of the DL. For this, cross-sectional images of the duct were identified and thresholded to generate a binary image with cilia cross-sections indicated by white pixels and "empty" lumen by black pixels. Then,

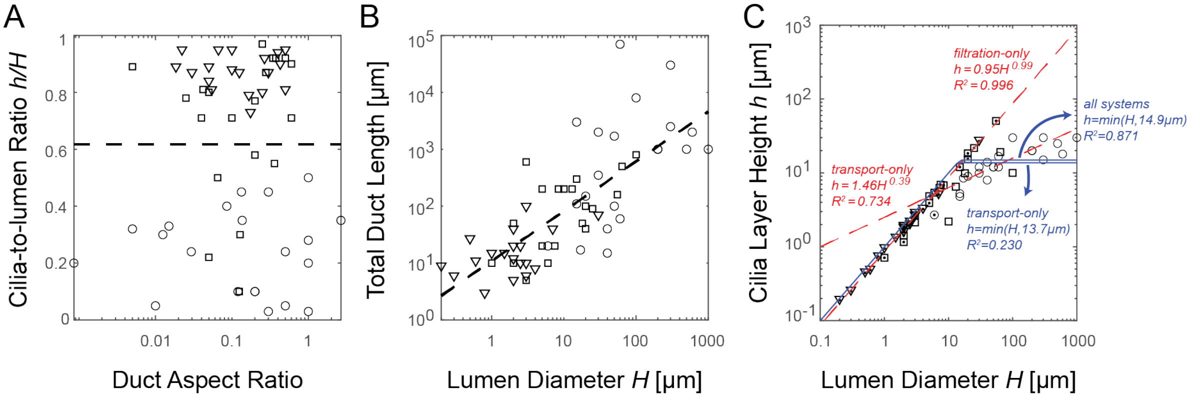


Figure S1: **Relationship of other morphometric parameters.** **A**, cilia-to-lumen ratio $h/H$ versus the duct aspect ratio – lumen diameter $H$ over total duct length with machine learning classification line overlayed. **B**, total duct length versus lumen diameter $H$ in log-log scale with a power law fit overlayed. **C**, different least-squares fits of cilia layer height $h$ as a function of lumen diameter $H$. The symbols follow that of Fig. 2: ○ for ciliated ducts with confirmed transport/mixing function, ▽ for filtration, and □ for unknown function.

$h/H$ was computed by dividing the cilia cross-sectional area (white pixels) by the total cross-sectional area of the DL (black and white pixels) and taking the square root.

**Animal phyla excluded from the analysis.** As indicated in Supplementary Table 1, phyla were excluded if literature suggested (1) absence of ducts with motile cilia (*e.g.*, Arthropoda, Orthonectida, Placozoa, Tardigrada, Xenacoelomorpha), (2) absence of any motile cilia (*e.g.*, Nematoda, Nematomorpha), or (3) ciliated duct geometries that could not be accurately described with parameters $h$ and $H$ (*e.g.*, Porifera).

**Classification using machine learning.** We conducted machine learning using the Decision Tree function with cross-validation in the Classification Learning App in Matlab (Mathworks, MA, USA). Five morphometric parameters (duct diameter and length, cilia length and orientation, and cilia-to-lumen ratio) were used to predict bulk transport versus filtration/valve function, where known (Supplementary Table 1). Results are shown in Fig. 2A and S1A.

**Total duct length and duct aspect ratio.** We visualize the relationship between cilia-to-lumen ratio $h/H$ and lumen diameter $H$ to the other recorded morphometric parameter total duct length. In Fig. S1)A, we see that adding duct length information in the form of a dimensionless duct aspect ratio (total duct length over lumen diameter) does not reveal any additional relationship other than what is revealed by the classification using $h/H$ alone discussed above. We believe this is because there is a simple, albeit noisy, correlation between duct length and lumen diameter, as shown by the power law fit line in Fig. S1B ($\log(L) = 0.87\log(H) + 1.04, R^2 = 0.568$).

**Least-square fitting of cilia layer height as a function of lumen diameter.** We show in Fig. S1C different least squares fits for cilia layer height $h$ as a function of lumen diameter $H$ using all surveyed systems. If there exists a universal limit on how long cilia can grow, $h$ should follow the functional form of $\min(h_{max}, H)$, since presumably cilia longer than lumen diameter will bend or buckle. Using all surveyed data together, this model give s fit with $h_{max} = 14.9$ [$\mu$m] with $R^2 = 0.8709$. However, this high coefficient of determination is misleading: If we only look at ducts with known bulk transport function, such a model can only produce a fit with $R^2 = 0.2303$ with $h_{max} = 13.7$ [$\mu$m]! However, using simple power-law fit commonly used for scaling-law analysis in biology, we can reach a fit of $R^2 = 0.7342$ with $h = 1.458H^0.393$ for bulk transport ducts and $R^2 = 0.9956$ with $h = 0.946H^0.985$ for ducts with known filtration functions (compared to $R^2 = 0.9804 using h = \min(27.0[\mu m], H)$). Therefore, we conclude that there exists a non-trivial relation that prompt cilia to growth to particular fraction of their ciliated ducts, and use our mathematical model to explore if there is a selection for pumping efficiency based on flow physics in addition to phylogenetic constraints. Trend lines discussed here are labeled in Fig. S1C.

#### S.2 Physics-based Modeling of Ciliary Ducts

We start by solving the Stokes' equation for $u_z^l$ in the central free lumen

$$\frac{\mu}{r}\frac{\partial}{\partial r}\left[r\frac{\partial u_z^l(r)}{\partial r}\right] = \frac{\partial p}{\partial z}, \qquad \text{for } 0 \leq r \leq a. \tag{1}$$

with finite velocity $u_z^l(0)$ at the center $r = 0$ and matching velocity at $r = a$,

$$u_z^l(a) = \varphi u_z^c(a), \tag{2}$$

where $\varphi$ is the isotropic fluid fraction inside the porous layer. Here, we assumed that the downstream velocity of the solids (cilia motion averaged over beat cycles) inside the porous ciliary layer is zero at leading order, similar to the infinitesimal analysis of channel flow with passive porosity in [4]. This assumption is also consistent with the notion that $f_c$ is the pressure gradient generated by the cycle-averaged ciliary forces pointing downstream in a confined pipe [5]. The analytical solution to Eq. (1) subject to the boundary condition in (2) is given by

$$u_z^l(r) = -\frac{\partial p}{\partial z}\frac{1}{4\mu}(a^2 - r^2) + \varphi u_z^c(a), \qquad \text{for } 0 \leq r \leq a. \tag{3}$$

The shear stress at the interface $r = a$ when approached from the free lumen side is given by

$$\sigma_{z,f}(a) = \mu\frac{\partial u_z^l}{\partial r}\bigg|_{r=a} = \frac{a}{2}\frac{\partial p}{\partial z}. \tag{4}$$

By continuity $\sigma_{z,p}(a) = \varphi\sigma_{z,f}(a)$ of shear stress $\sigma_{z,p}$ and $\sigma_{z,f}$ at $r = a$ [4, 6], we get

$$\mu\frac{\partial u_z^c}{\partial r}\bigg|_{r=a} = \varphi\frac{a}{2}\frac{\partial p}{\partial z}. \tag{5}$$

The Brinkman equation in the ciliary layer is given by

$$\frac{\mu}{r}\frac{\partial}{\partial r}\left[r\frac{\partial u_z^c(r)}{\partial r}\right] = \varphi\frac{\partial p}{\partial z} + K_c u_z^c(r) + f_c, \qquad \text{for } a \leq r \leq R, \tag{6}$$

subject to (5) and the no-slip boundary condition at the duct wall $u_z^c(R) = 0$. The analytical solution of (6) is given by

$$
\begin{aligned}
u_z^c(r) = &\left[2\left(\mathrm{I}_1(\hat{a})\mathrm{K}_0(\hat{R}) + \mathrm{K}_1(\hat{a})\mathrm{I}_0(\hat{R})\right)\right]^{-1} \\
&\left[2(\tilde{f} - \delta p)\left[\mathrm{I}_1(\hat{a})\left(\mathrm{K}_0(\hat{R}) - \mathrm{K}_0(\hat{r})\right) + \mathrm{K}_1(\hat{a})\left(\mathrm{I}_0(\hat{R}) - \mathrm{I}_0(\hat{r})\right)\right]\right. \\
&\left. - \hat{a}\delta p\left[\mathrm{I}_0(\hat{R})\mathrm{K}_0(\hat{r}) - \mathrm{I}_0(\hat{r})\mathrm{K}_0(\hat{R})\right]\right], \qquad \text{for } a \le r \le R,
\end{aligned}
\tag{7}
$$

where $\hat{(\cdot)} = \sqrt{\frac{K_c}{\mu}}(\cdot)$, $\delta p = \frac{\varphi}{K_c}\frac{\partial p}{\partial z}$, $\tilde{f} = \frac{f_c}{K_c}$, and $\mathrm{I}_{(\cdot)}, \mathrm{K}_{(\cdot)}$ are modified Bessel functions of order $(\cdot)$.

Note that since Equation (7) is linear in $\delta p$ and $\tilde{f}$, we only need to compute solution $U_c = u_z(\tilde{f} = 1, \delta p = 0)$ and $U_p = u_z(\tilde{f} = 0, \delta p = 1)$ and obtain $u_z(\tilde{f}, \delta p) = \tilde{f}U_c + \delta p U_p$.

It is interesting to note that for $\partial p/\partial z = 0$, $u_z$ becomes independent of $a$ as $R \to \infty$. In other words, at zero adverse pressure, the cilia driven flow speed becomes independent of $h/H$ for large $H$. This trend dictates the existence of an optimal efficiency at small adverse pressure because the amount of active material increases with higher $h/H$.

### S.3 Effective parameter selection

**Setting lower limit of the cilia solid fraction $\varphi_c$ in the ciliary carpet layer.** We seek an estimate of the cilia solid fraction $\varphi_c$ within the ciliary carpet layer. Since the cilium diameter and inter-cilium spacing are roughly equal (about 0.2 $\mu$m), in a uniformly-covered ciliated tissue, the expected cilia density should be about $\varphi_c = 0.5^2 \equiv 25\%$. However, many ciliary carpets contain both ciliated and non-ciliated cells, such that the tissue-level coverage fraction of ciliated cells is much less than 100%. In mouse trachea, the ciliated cell coverage can be as low as 37% [7], and amphibian ciliated skin exhibits a 50% ciliated cell coverage [8]. Additionally, individual ciliated cells can have partial ciliation, such as in ependymal epithelia in mammalian brain ventricles where the ciliated area overall approaches only 32% of total surface [9, Fig. 3]. Thus, the overall cilia density, accounting for this heterogeneity in surface coverage is far less than 25%. For example, a tissue with 30%–50% coverage fraction, at 25% cilia density within the covered patches would have an overall cilia density of about 7.5%–12.5%. Thus, we used $\varphi_{c,min} = 10\%$ as a lower bound for our cilia solid fraction (see Fig. 3E of the main text). Together with the ansatz $\varphi_c = h/H$ for observed ciliary flame systems, we use a softplus function $0.1 \log(1 + \exp(10(h/H - \varphi_{c,min}))) + \varphi_{c,min}$ to represent $\varphi_c$ for all values of $h/H$ regardless of the ciliated duct type.

**Setting dimensional scale of the active force density $f$.** To calibrate the correct order of magnitude for $f$, we used measurements of pumping performance in the ciliary flames that connect the peritoneal cavity to the vasculature and filter lymphatic in the toad *Bufo marinus* and *Bufo bufo* [10, 11]. This filtration system consists of 600–800 ciliated flames. Flames exhibit circular and elliptical apertures between 7–40 $\mu$m in diameter [10]. Although openings as large as 100 $\mu$m were reported, many larger flames appeared to have flap covers that would reduce their “hydrodynamic" diameter [10, Fig.4]. No clear length was visible or reported in [10], but similar ciliary flames have lengths of at least 30 $\mu$m [11, Fig. 7c]. By measuring the pressure difference between the peritoneal and the blood compartment, the authors of [10] reported a maximum flow rate of 0.5±0.03 ml/hr, a flow rate of 0.17 ml/hr at about 200 Pa, and a maximum pressure of 310 $\pm$ 20 Pa (see [10, Fig. 12]). A summary of these measurements is given in Table S1. As a side note, in our survey of ciliated organs in Fig. 2 of the main text, we used a specific *Bufo marinus* funnel of $H = 7$ $\mu$m and $h/H = 0.73$ shown in [10, Fig.11] due to its clear visualization of cilia density. To calibrate our model and estimate the force density, we considered in the context of our model the specific example of a ciliated flame of average lumen diameter 15 $\mu$m and length 100 $\mu$m. We used the measured cilia-to-lumen ratio of $h/H = 0.73$

and the parameter values $\kappa/\mu = 1\ \mu\text{m}^{-2}$ and $\mu = 10^{-3}$ Pa·s corresponding to the viscosity of water. Our model predicted that this funnel matches the maximum pressure of about 300 Pa observed in [10] if we set the active force density to $f = 15$ pN/$\mu$m$^3$. This value is well within the mechanical capability of cilia averaged over a stroke cycle, given that the internal motors of a single respiratory cilium can generate about $60$ pN of force during its effective stroke at the tip, implying a force density of nearly 200 pN/$\mu$m$^3$ for a $200$ nm wide, $10\ \mu$m long cilium [12]. To further verify the validity of the estimate of $f = 15$ pN/$\mu$m$^3$ for flames of dimensions $100\ \mu\text{m} \times 15\ \mu\text{m}$ and $h/H = 0.73$, we calculated that a total of 800 such flames produce a flow rate of about 0.16 ml/hr against 200 Pa of pressure with a maximum attainable flow rate of about 0.46 ml/hr, which are very close to the experimentally reported flow rates. A comparison of experimental data and our model predictions are available in Table. S1. As noted in § S.2, because the final flow field solution depends linearly on the ratio $f_c/K_c$, any choice of force density scale $f$ indepdendent of $h/H$ and $H$ will not alter the optimality trends computed by our method. Therefore, it is also sensible to consider flow speed/rate derived using our model with other values of $f$, where the effective forward pressure gradient generated by cilia activity is known to be different, possibly due to changes in unmodeled, microscopic details of the cilia coordination or beat waveforms.

**Effect of changing the effective Brinkman coefficient $K_c = \kappa\varphi_c$.** Since the Brinkman drag coefficient is fundamentally empirical for geometrically complex systems [15, 16], we assumed for simplicity that $\kappa$ is of the same order as the fluid viscosity throughout the main text of our study. However, since $\sqrt{\mu/\kappa}$ can also be interpreted as an effective pore size for the porous layer [17], it is interesting to see how changing $\kappa$ would affect the results of our optimization algorithm (Supplementary Algorithm S1). In Fig. S2, we show that increasing the relative value of $\kappa/\mu$ (decreasing the effective pore size) decreases the lumen diameters that maximize the

| Filter Type (organism) | Sustainable Pressure [kPa] | Total Filtration Rate [mL/hr] | Unit Size (L x Ø) [μm] | Unit Volume [μL] | Number of Units Required | Total Volume [μL] |
|---|---|---|---|---|---|---|
| **Blood-pressure-based Filter** (Human kidney glomeruli) | 5 − 10 | 7500 | N/A | 0.006 | 1,000,000 | 6,000 |
| **Cilia-powered Filter** (*B. marinus* peritoneal flame) | 0.29 − 0.33 | 0.17 @ 200 Pa | unknown x 7 − 40 | unknown | 600 − 800 | unknown |
| **Cilia-powered Filter** (simulated) | 0.3 | 0.16 @ 200 Pa | 100 x 15 | 0.00002 | 800 | 0.01 |
| **Larvacean Ciliated Funnel** (*Bathochordaeus sp.*) | 0.2 | 0.4 @ 100 Pa | 160 x 55 | 0.0004 | N/A | N/A |
| **Cilia-powered "human kidney"** | 10 | 7500 @ 5 kPa | 8000 x 55 | 0.02 | 20,000,000 | 400,000 |

Table S1: **Pumping characteristics of observed and simulated filtration systems.** We list the structural and functional parameters found in literature (black) and derived based on our model (blue). Data for human kidney glomeruli are approximated from [13, 14]. Data from the cilia powered peritoneal filtration in *Bufo marinus* are based on [10]. In all simulations, we used $f = 15$ pN/$\mu$m$^3$, $\kappa/\mu = 1$ $\mu$m$^{-2}$, and $\mu = 10^{-3}$ Pa·s.

pumping efficiency $E$. Importantly, all values of $\kappa$ produce the same relationship between the value of adverse pressure gradient d$p$/d$z$ and cilia-to-lumen ratio $h/H$, implying that different values of $\kappa$ would only move the most efficient designs (solid white lines in Fig. 4B or solid black lines in Fig. 4C-D) laterally in the $(H, h/H)$ morphospace;see Fig. S2B. This effect adds another reason why some of the biological ciliated ducts deviates laterally from the most efficient line obtained for the value $\kappa/\mu = 1[\mu m^{-2}]$ (Fig. S3A); they could be more efficient for a different value of effective pore size. Lastly, because predicted $h/H$ remain invariant for all choices of $\kappa$, the log-log slope between the cilia layer height $h$ and lumen diameter $H$ for the most efficient systems is also invariant with respect to $\kappa$; we show the computed result for $\kappa/\mu = 1[\mu m^{-2}]$ in Fig. S3B. Note that for most lumen diameter values, the predicted slope is comparable to the power law fits of $0.393 \approx 0.4$ shown in Figure 2 and 4 as red dashed lines.

**Algorithm S1** Evaluating performance of duct designs on the morphospace $(H, h/H)$ and identifying optimal duct designs that maximize efficiency $E$ under adverse pressure.

**Require:** list of ciliated duct morphological parameters:
$(H, h/H)$ // morphospace formed by lumen diameters and cilia-to-lumen ratios
$\mu, f, \varphi_c(H, h/H)$ // fix viscosity, active force scale and forms for cilia solid fraction.
**for** $\kappa/\mu = 0.1, 1, 10$ **do**
Pre-compute the following and store each as a matrix over the morphospace $(H, h/H)$:
$U_c \equiv U_z(H, h/H)$ for $f = 1, \mathrm{d}p/\mathrm{d}z = 0$ // flow velocity due to unit cilia activity
$U_p \equiv U_z(H, h/H)$ for $f = 0, \mathrm{d}p/\mathrm{d}z = 1$ // flow driven by unit adverse pressure
$A_c \equiv \frac{1}{4}\pi\varphi_c H^2(1-(1-h/H)^2)$ // total ciliated area per cross section
Compute maximum possible pressure and flow rate generation over $(H, h/H)$:
$Q^* \equiv \frac{1}{4}\pi H^2 U_c$ // Maximum flow rate occurs when adverse pressure is absent.
$\mathrm{d}p/\mathrm{d}z^* \equiv -f \cdot U_c/U_p$ // Maximum pressure is generated when net flow is zero.
Initialize null relative efficiency score over the entire morphospace $(H, h/H)$:
$E_{\mathrm{rel}} \equiv$ **zeros**$(H, h/H)$
**for** $\mathrm{d}p/\mathrm{d}z = 10^{-10}, \ldots, 10^2$[Pa/$\mu$m] **do**
$U = f \cdot U_c + \mathrm{d}p/\mathrm{d}z \cdot U_p$ // Compute average flow speed at given adverse pressure.
**if max**$(U) < 0$ **then**
**break** // Quit if no design parameters can pump at this $\mathrm{d}p/\mathrm{d}z$.
$E = Q/A_c = \frac{1}{4}\pi H^2 U/A_c$ // Compute efficiency at given adverse pressure.
Identify optimal designs at given $\mathrm{d}p/\mathrm{d}z$:
$[E^\dagger, (H, h/H)^\dagger] =$ **max**$(E)$ // Find design parameters that maximize efficiency
$E_{\mathrm{rel}} =$ **max**$(E_{\mathrm{rel}}, E/E^\dagger)$ // Accumulate relative efficiency score
**return** $(H, h/H)^\dagger$ // Return optimal design parameters as function of $\mathrm{d}p/\mathrm{d}z$.
**return** $Q^*, \mathrm{d}p/\mathrm{d}z^*, E_{\mathrm{rel}}$ // Return quantities as function of $\kappa/\mu$.

#### S.4 Quantitative comparison of excretory organs

Flame cells serve as a basic filtration unit in many mm-scale organisms, but are markedly absent in larger organisms, (except during some stages of development) [18]. Filtration in larger organisms, the human kidney for example, leverages the blood pressure generated by a muscular heart. We propose that the lack of filtration by ciliary flames in larger animals might be (partly) due to the unfavorable scaling of pressure and flow generation in flame-based filtration compared to blood pressure-based filtration, as the following example illustrates. In humans, the filtration pressure provided by the capillary blood pressure is on the order of $5-10$ kPa [13].

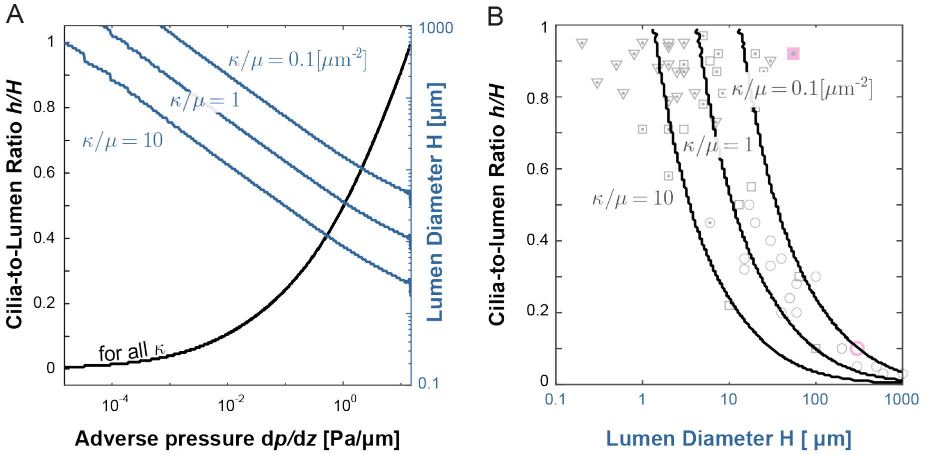


Figure S2: **Effect of Brinkman coefficient $\kappa$ on morphology of most efficient designs. A.** Result of our optimization (Algorithm S1) for different values of $\kappa$ while maintaining all other parameters fixed. The most efficient designs have smaller lumen diameter $H$ at a higher hydraulic resistance coefficient $\kappa/\mu$, but the optimal cilia-to-lumen ratio $h/H$ do not change. **B.** In morphospace, the most efficient designs shift only horizontally for different values of $\kappa$. This provides an additional explanation for the lateral spread of the biological data points. Here $\mu = 10^{-3}$ [Pa$\cdot$ s] and $f = 15$ [pN/$\mu$ m$^3$].

The average filtration rate of the kidney is ca. 180 liters of blood per day, which, considering nearly 1,000,000 filtration units (glomeruli) [14], indicates a filtration rate of 7.5 $\mu$l/hr per glomerulus. Given an average glomerular volume of $6 \cdot 10^6$ $\mu$m$^3$, the total space taken up by the filtration units is ca. 6 ml [14]. The larvacean's ciliated funnel, the largest ciliary pump found in our survey, can efficiently pump 0.4 $\mu$l/hr of fluid against 0.6 Pa/$\mu$m of pressure gradient (Fig. 4A). This means that the parallel operation of ca. 20 such funnels, each with more than 8 mm in length, can produce comparable a flow rate (20×0.4 = 8 $\mu$l/hr) at the minimum 5 kPa pressure drop observed across the glomerular filtration barrier. Assuming biologically feasible, such funnel would be able to sustain a maximum pressure at nearly 10 kPa without flow. Using the lumen diameter $H = 55$ $\mu$m as a lower bound, the resulting tissue volume that replaces a single glomerulus would measure 0.02 $\mu$l, which leads to about $0.4$ liters if these funnels were to replace the operation of all 6 ml of glomerular volume in an entire kidney. This example demonstrates the unfavorable scaling of cilia-based filtration to larger body sizes and

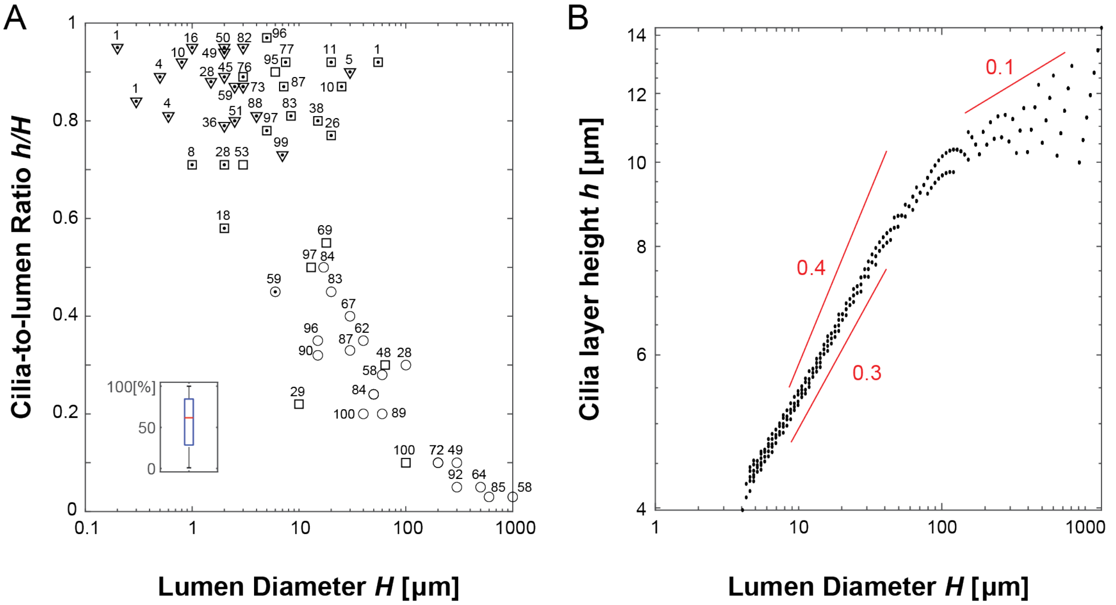


Figure S3: **Predicted efficiency assuming** $\kappa/\mu = 1[\mu m^2]$**. A.** We show the maximum relative efficiency produced by our optimization algorithm at the exact point of morphospace associated with each reported data points. From the box plot (inset), we see that most of the system lie reasonably close to the most efficient line predicted by our hydrodynamic model without parameter fine-tuning. **B.** The log-log slope of the computed most efficient systems is between 0.3 and 0.4 for lumen diameter below 100 [$\mu$m], and drops to around 0.1 for larger lumen diameters. These slope is invariant with respect to the specific parameter choices. Here $\mu = 10^{-3}$ [Pa· s] and $f = 15$ [pN/$\mu$ m$^3$].

their higher filtration rate demands.
